# Cognitive reappraisal of food and emotion cues involves common and unique neural contributions

**DOI:** 10.64898/2026.03.09.710570

**Authors:** Julia M. Laing-Young, Cary R. Savage, Cara Tomaso, Maital Neta, Timothy D. Nelson, Douglas H. Schultz

## Abstract

Obesity is a growing public health concern with more than 40% of adults meeting criteria for obesity in the United States. Although many treatments seek to lower individual’s weight, few treatments have focused on cognitive strategies to change the way individuals think about food, therefore, decreasing consumption of non-nutrient-dense foods. Cognitive reappraisal is one strategy that involves changing the way one thinks about a situation and can be used to downregulate responses to those stimuli. Leveraging this intuitive, cost-effective strategy to decrease one’s desire to eat unhealthy food and therefore, decrease overeating, could improve physical and mental health. The present study identified brain regions that are differentially activated when using cognitive reappraisal to downregulate responses to food (FR) versus when using the same strategy to downregulate negative emotions (ER). We collected functional magnetic resonance imaging (fMRI) data in 63 undergraduate students while participants completed both tasks. There was increased reappraisal-related activation in widespread regions across both tasks, including in expected subcortical (i.e., striatum) and cortical areas (i.e., visual, frontoparietal). We also found domain-specific activity, with greater insula activation in the FR than the ER task and greater hippocampal activation in the ER than the FR task. These results reveal domain-general and domain-specific effects of cognitive reappraisal in FR and ER tasks that inform future work examining eating behavior. Taken together, a better explication of the overlapping and discrete processes of food regulation, as it compares to other applications of this regulatory strategy can inform new intervention targets.

## Introduction

Obesity is a growing public health concern. In the United States, more than 40% of adults meet criteria for clinical obesity (body mass index [BMI] of 30 or higher), and an additional 30% of adults are classified as overweight (National Center for Health Statistics, Centers for Disease Control and Prevention, 2024). Obesity is a complex and difficult to treat chronic disease associated with significant health consequences, including diabetes, cardiovascular disease, and even mortality (Koliaki et al., 2023). The obesity epidemic is further fueled by the consequences of economic, social, and technological advances in recent decades which promote increased calorie consumption and reduced physical activity, and place increased demands on self-regulation. For example, palatable foods with high caloric density are easily available in fast food restaurants and prepackaged items (Pi-Sunyer, 2002; Catenacci et al., 2009), the increase in popularity of food delivery services (e.g., DoorDash, UberEats), and Americans are more likely than Europeans to use automobiles for their primary form of transportation, resulting in less walking (Buehler & Hamre, 2014; Buehler & Pucher, 2023). To decrease obesity prevalence, interventions are needed at the individual level, for example, by decreasing individual-level food cravings (i.e., the desire to eat high caloric density, readily available foods).

The ability to regulate food cravings is associated with reduced risk for obesity through lower caloric consumption (Potenza & Grolo, 2014; Sun & Kober, 2021). These reductions in caloric consumption demonstrate the importance of food regulation for physical wellbeing. Cognitive reappraisal is a key regulation strategy used to down-regulate negative emotions and food cravings and is defined as a cognitive strategy that involves deliberately changing how one interprets the meaning or salience of a given stimulus (McRae, Ciesielski, & Gross, 2012a). In the context of negative emotions, cognitive reappraisal may be used to help an individual find less negative or even identify positive interpretations of an emotional stimulus (e.g., viewing a bruised body as a scene from a movie). In the context of food regulation, the individual might focus on the health consequences of consuming unhealthy foods to decrease their desire for a meal or snack.

Recent functional magnetic resonance imaging (fMRI) research has investigated the neural processes involved in regulatory behavior, specifically food regulation (FR) and emotion regulation (ER). These studies have identified several brain regions that are involved when participants reappraise food stimuli relative to when they passively view food stimuli or are allowed to crave them. This core set of brain regions, including the anterior insula, inferior and middle frontal gyrus, supplementary motor cortex and parietal cortex – components of the cingulo-opercular and frontoparietal networks (Hausman et al., 2021) – have been broadly implicated in cognitive control (Vincent et al., 2008; Zanto et al., 2013). Other work has also linked these regions with dietary self-control (Han et al., 2018). Given that, it is not necessarily clear the extent to which these effects apply to regulation abilities in general rather than being specific to food regulation.

The use of cognitive reappraisal to downregulate responses to negative emotional images is associated with activation in various regions of the prefrontal cortex (PFC; e.g., dorsolateral, ventrolateral, and dorsomedial PFC, the pre-supplemental motor area, and the anterior cingulate cortex; Chambers et al., 2009; Buhle et al., 2014), which are also regions of the cingulo-opercular and frontoparietal networks (Friedman & Robbins, 2021; Sokolowski et al., 2021). A meta-analysis (Zilverstand et al., 2017) found clinical populations consistently demonstrate reduced recruitment of the dorsolateral and ventrolateral PFC when downregulating negative emotions, suggesting there may be a core deficit in this reappraisal process. As noted in the context of food regulation, these brain regions are broadly implicated in domain-general cognitive control and may not be specific to emotion regulation.

Despite it being unclear whether these regulatory processes involve overlapping or distinct neural processes, food dysregulation and emotion dysregulation are similar in that they are both associated with clinical presentations. The inability to control food cravings is associated with overeating and obesity (Giuliani et al., 2013), and the inability to regulate negative emotions is a core component of mood disorders (Farb et al., 2012). Further, food dysregulation is a common feature of depression, and individuals with difficulty regulating their emotions may use food (i.e., binge-eating or food restriction) as an attempt to cope (Dingermans, Danner, & Parks, 2017). This example illustrates a potential mechanistic link between food and emotion dysregulation. Although FR and ER have been studied independently (Ochsner & Gross, 2005; Pruessner et al., 2020; Wardle, 1988; Kleiman et al., 2016; Davidson & Jones, 2019), minimal research has directly compared their underlying neural mechanisms using a within subjects design. A recent between-subjects meta-analysis (Gerosa et al., 2024) found a large overlap between brain regions involved in downregulating food cravings and emotional stimuli (e.g., dorsolateral and ventrolateral PFC), as described above. There were also domain-specific effects for food cravings, notably activation in the right insula and left inferior frontal gyrus. However, differences in task design across ER and FR limited the interpretation of these comparisons.

To address this gap in the literature, we examined FR and ER using a similar task design in the same set of participants. The aim of this study was to identify the brain regions associated with cognitive reappraisal during both FR and ER, referred to as domain-general hereafter, and brain regions uniquely associated with only one of these tasks, referred to as domain-specific. We hypothesized that domain-general cognitive reappraisal would recruit activation in cognitive control regions (e.g., dorsolateral PFC) that are affiliated with fronto-parietal and cingulo-opercular networks. Additionally, we hypothesized domain-specific effects in reward-related brain regions, such as the insula, striatum, and orbitofrontal cortex, for the FR task relative to the ER task. Identifying overlapping and unique mechanisms for these regulatory tasks has the potential to inform future intervention strategies, which could in turn improve mental and physical health throughout the lifespan. For example, if FR and ER have distinct mechanisms, this research could provide insight into food dysregulation and emotion dysregulation requiring domain-specific interventions to target health outcomes.

## Methods

### Participants

Sixty-eight undergraduate students were enrolled from the University of Nebraska-Lincoln’s psychology research participant pool and received course credit for participation. Eligibility criteria included being 18 or 19 years old, right-handed, and no MRI contraindications (e.g., metallic implants, claustrophobia, pregnancy). Of the 68 participants enrolled that completed the scanning procedures, 63 were included in the analyses (*M_age_* = 19.13*; M_BMI_* = 24.13; 75% female; 85.9% white, 6.3% Asian American, 4.7% Black, 3.1% did not report; 10.7% Hispanic/Latinx). Neuroimaging data for two participants were unusable due to high motion (>33% of datapoints during tasks were censored), and three participants were excluded due to incomplete data (i.e., missing behavioral ratings on tasks).

### Procedures

Participants who met inclusion criteria attended a multi-hour laboratory visit at the University of Nebraska-Lincoln’s Center for Brain, Biology, and Behavior (CB3). At the start of the session, participants confirmed that they had not consumed any food or caffeinated beverages during the four hours prior to their scheduled appointment. Participants then completed the informed consent process with a trained research assistant, as well as having their height and weight measured. Prior to starting the fMRI session, participants were trained on fMRI tasks to ensure they were appropriately using reappraisal strategies. All study procedures were approved by the Institutional Review Board at the University of Nebraska-Lincoln (UNL).

### fMRI Tasks

#### Food Regulation Task

Participants completed a food regulation task adapted from Yokum and Stice (2013), in which they were shown 48 appetitive food images (Figure 1). The task was presented using E-Prime software (version 2.0) (Psychology Software Tools, Inc.). Images were selected from the Open Library of Affective Foods (OLAF; Miccoli et al., 2016), supplemented with images of high calorie foods used in previously published work (Ness et al., 2014). All images were rated ≥6 on a 1-9 craving scale in this previous online pilot study. During the *IMAGINE* condition (24 trials), which was cued with the word “LOOK,” participants were instructed to imagine themselves eating the food, including what the food might smell and taste like. During the *REAPPRAISAL* condition (24 trials), which was cued with the word “DECREASE,” participants were trained to regulate their responses by thinking about the long-term health consequences of eating the food (e.g., “Eating fried food may lead to future unwanted weight gain”). Participants were encouraged to consider health consequences that are most relevant to their personal health goals. This strategy was selected because of its face validity with appetitive executive control abilities and because it is very similar to other effective strategies used to decrease appetitive responses in earlier work (Yokum & Stice, 2013).

**Figure 1:**
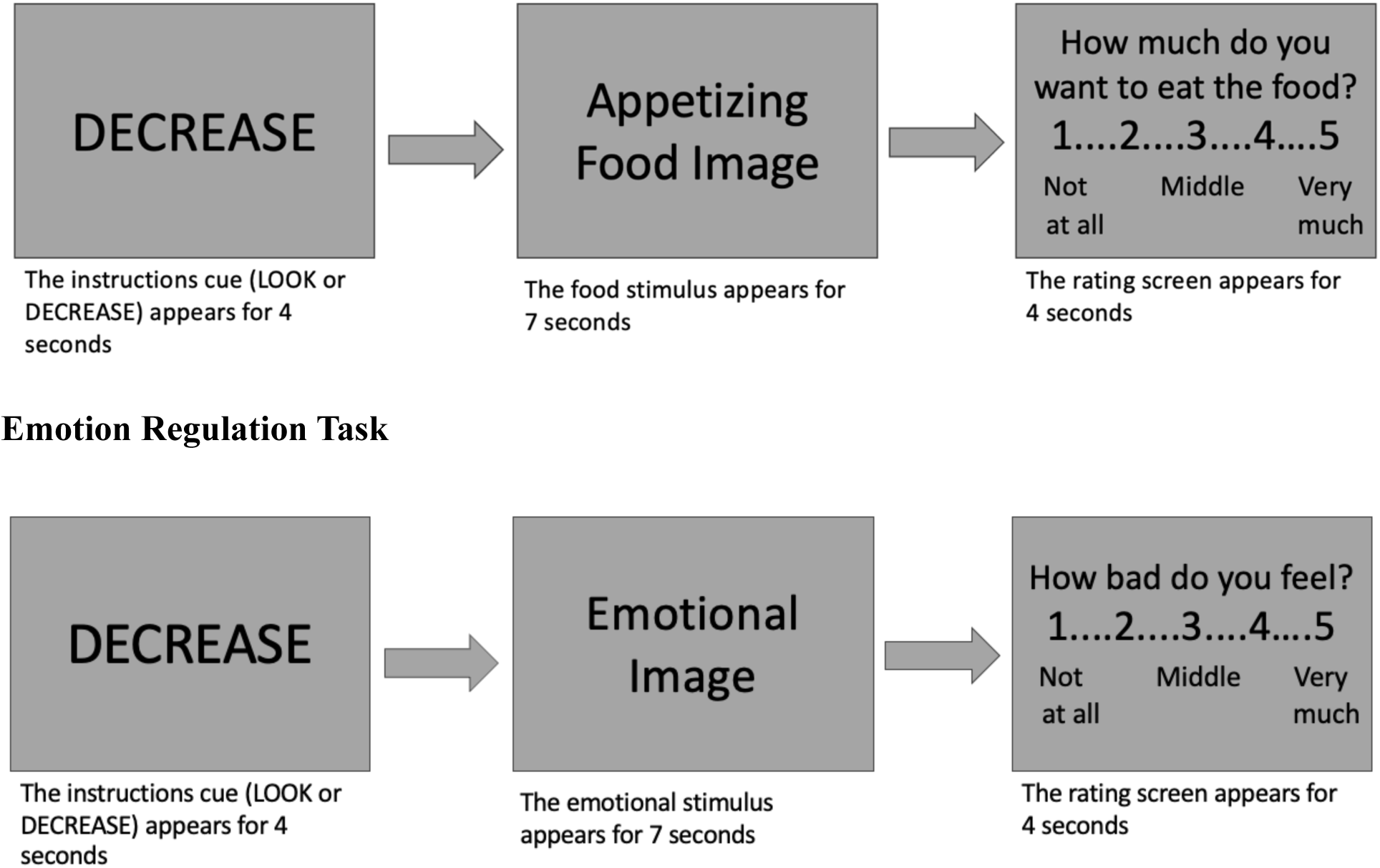
Food Regulation Task. (adapted from Yokum and Stice (2013)): During the imagine condition (cued with LOOK), participants were instructed to imagine themselves eating the food that appeared in the subsequent image. During the reappraisal condition (cued with DECREASE), participants were instructed to think about the long-term negative health consequences of eating the food that appeared in the subsequent image. After the image disappeared, they rated how much they wanted to eat the food on a 1 (not at all) to 5 (very much) scale. **Emotion Regulation Task** (adapted from McRae et al., 2012): When the “LOOK” cue appeared, participants were instructed to participants were instructed to respond naturally and allow whatever feelings may arise. During the reappraisal condition (cued with DECREASE), participants were instructed to think about three types of re-interpretation: It is not real (i.e., it’s just a scene from a movie, they are pretending), things will improve with time, and things are not as they appear to be. After the image disappeared, they rated how much they wanted to eat the food on a 1 (not at all) to 5 (very much) scale.

During training, participants verbalized their strategies to the *IMAGINE* condition and to the *REAPPRAISAL* condition to ensure they could quickly and effectively think of negative health consequences without using a non-cognitive strategy (e.g., attention deployment strategies such as looking away from the image). If participants’ responses while training revealed other strategies, the research assistant provided corrective feedback until they could successfully use the reappraisal strategy. For each trial, the condition instruction (i.e., “LOOK” or “DECREASE”) appeared at the center of the screen for 4 seconds. Then a food image was presented for 7 seconds, followed by a prompt that appeared for 4 seconds asking participants to press a button rating their desire for the food on a 5-point scale (1 = not at all to 5 = very much). A variable intertrial interval (2-4 seconds) separated each trial, during which time participants were instructed to relax and clear their minds. The entire task lasted 12 minutes, and the order in which the trials were presented was pseudo-randomized such that no more than two sequential trials were of the same condition. Conditions were also counterbalanced, and “LOOK” and “DECREASE” stimuli were randomly assigned for each participant.

#### Emotion Regulation Task

Participants completed an emotion regulation task adapted from McRae et al., 2012 (Figure 1) in which they were shown a total of 60 unique images. The task was presented with E-Prime software (version 2.0) (Psychology Software Tools, Inc.). Images, which were selected from the International Affective Picture System (IAPS; Lang et al., 2008), included 40 negative and highly arousing images taken from prior research (Neta et al., 2022). Each trial began with an instruction screen lasting 4 seconds with either “LOOK” or “DECREASE” written on a green or blue background, respectively. This was followed by a presentation of an image for 7 seconds against the same-colored background as the instruction screen. For the “LOOK” instruction trials, half of the images had a negative valence (“Look Negative”, 20 trials) and half of the images had neutral valence (“Look Neutral”, 20 trials); participants were instructed to respond naturally and allow whatever feelings may arise. For the “DECREASE” instruction trials (*REAPPRAISE*; 20 trials), all the images were negative, and participants were instructed to use cognitive reappraisal to reinterpret the content to feel less negative (see more below). Next, a rating screen appeared for 4 seconds and participants were asked to indicate the degree of negative emotion they felt at the end of the image presentation (i.e., after reappraisal or look.) Participants were asked, “How bad do you feel?” on a 5-point scale (1 = not bad at all; 5 = very bad). A variable intertrial interval (2-4 seconds) separated each trial, during which time participants were instructed to relax and clear their minds. The entire task lasted approximately 16 minutes, and as in the FR task, trials were presented pseudo-randomly, and the negative images that were assigned to “LOOK” or “DECREASE” were counterbalanced across participants.

Prior to task onset, participants were given instructions and shown example images not used during the task itself. Participants completed a set of three practice trials (two “DECREASE” and one “LOOK” trial). For the “DECREASE” trials, the experimenter suggested three types of re-interpretations: It is not real (i.e., it’s just a scene from a movie, they are pretending), things will improve with time, and things are not as they appear to be. During the training, participants were asked to explain how they reappraised each scene. If participant responses during training revealed that they were not using a reappraisal strategy, the experimenter offered corrective feedback and re-directed the participant to use one of the three re-interpretation strategies provided above.

### fMRI image acquisition

fMRI data were collected with a 3.0-T Skyra Siemens scanner with a 64-channel head coil. Prior to functional imaging, high-resolution anatomical scans were acquired, T1 weighted multi-echo magnetization prepared rapid gradient echo sequence (MP-RAGE; TR = 2530 msec, TE1 = 1.69 msec, TE2 = 3.55 msec, TE3 = 5.41 msec, TE4 = 7.27 msec, matrix size = 256 x 256 mm^2^, field of view = 256 mm, slice thickness = 1mm, number of sliced = 176, and TA = 6.03 min). Functional scans used a multi-band factor = 8, accelerated echo planar imaging (EPI) sequence with the following parameters: repetition time (TR) = 1000 msec, time of echo (TE) = 29.80 msec, flip angel, 60 degrees, matrix size = 210 x 210 mm^2^, field of view = 210 mm, slice thickness = 2.5 mm, number of sliced = 51, and voxel size = 2.5 mm^3^. Throughout the imaging protocol, participants’ heads were secured with foam pads and participants wore foam earplugs. Participants provided ratings for their food desire (FR task) and for how badly they felt after viewing the images (ER task) using a Celeritas Fiber Optic Five-Button Response System placed on their right side (Psychological Software Tools, Inc.). Lastly, movement during the tasks was monitored using Framewise Integrated Real-Time MRI Monitoring (FIRMM) software (Dosenbach et al., 2017).

### MRI preprocessing

Functional imaging preprocessing was performed using Analysis of Functional Neuroimages (AFNI; Cox, 1996). Anatomical images were segmented using FreeSurfer (Fischl et al., 2004; Desikan et al., 2006). Processing was completed using afni_proc.py and involved despiking, slice time correction, standardization of anatomical images to Montreal Neurological Institute (MNI) template (152_2009), volume registration to the minimum outlier fraction, spatial smoothing using a 4 mm full width at half maximum Gaussian filter, the time course of each voxel scaled to have a mean of 100, and a general linear model (GLM) using the six motion estimates from volume registration as regressors of no interest. We also used up to third-order polynominals to model baseline and drift. Pairs of volumes where the Euclidean norm of the motion derivatives exceeded 0.3 were censored and eliminated from further analysis.

To compare the effects of the regulate and look/imagine strategies, blood oxygenation level-dependent (BOLD) responses for each condition were contrasted. We used the “BLOCK” basis function to model response functions at the start of each task trial. Specifically, the “BLOCK” basis function had a duration of 4 seconds for the instruction prompt (i.e., LOOK or DECREASE cue) and 7 seconds for the images.

### Primary data analysis

AFNI’s 3dMVM (Chen et al., 2014) was used to conduct a multivariate modeling analysis with task (FR, ER) and condition (regulate vs. imagine/look) as within-subject variables. The spatial smoothness of participants’ residuals was estimated with the “-acf” option (Cox et al., 2017). Mean values for the sample were 0.7151, 2.0648, 6.2374, respectively. Next, a cluster-based approach for multiple comparisons correction was used to identify which regions showed a significant main effect or interaction. Specifically, AFNI’s 3dClustSim was used to generate ten thousand random maps using the smoothness parameters and thresholded at a voxel-wise *p* < 0.001. The largest surviving cluster from each of these simulations was recorded, and this distribution was used to estimate the probability of a false positive. Based on these estimations, we selected a voxel-wise threshold of *p* < 0.001 and a minimum cluster size of eight contiguous voxels, resulting in a corrected two-tailed alpha of *p* < 0.05. This resulted in a map for the main effect for task (FR vs. ER), the main effect for condition (regulate vs. imagine/look), and the interaction.

Finally, post-hoc tests were conducted to interrogate the interaction effects. Using the previously identified interaction clusters, the means from the look condition were subtracted from the regulate condition for each task. We categorized each cluster as showing a greater differential activation FR or ER by comparing the absolute value of the ‘look – regulate’ for each task and assigning it to a respective task based on which mean was furthest away from zero. The absolute value of ‘look – regulate’ activations for each task were also correlated with a difference score of ‘look – regulate’ behavioral ratings (i.e., regulation success).

## Results

### Behavioral results

Participants were instructed, on some trials, to use cognitive reappraisal while viewing food or emotional stimuli, and on other trials, to look at the images allowing themselves to feel what they would naturally feel. As expected, ratings of food cravings on decrease trials were lower than look trials in the FR task (*t* = 14.23, *p* = <0.001, Figure 2) and ratings of negative affect on decrease trials were lower than look trials in the ER task (*t =* 14.10*, p* = <0.001, Figure 2).

**Figure 2:**
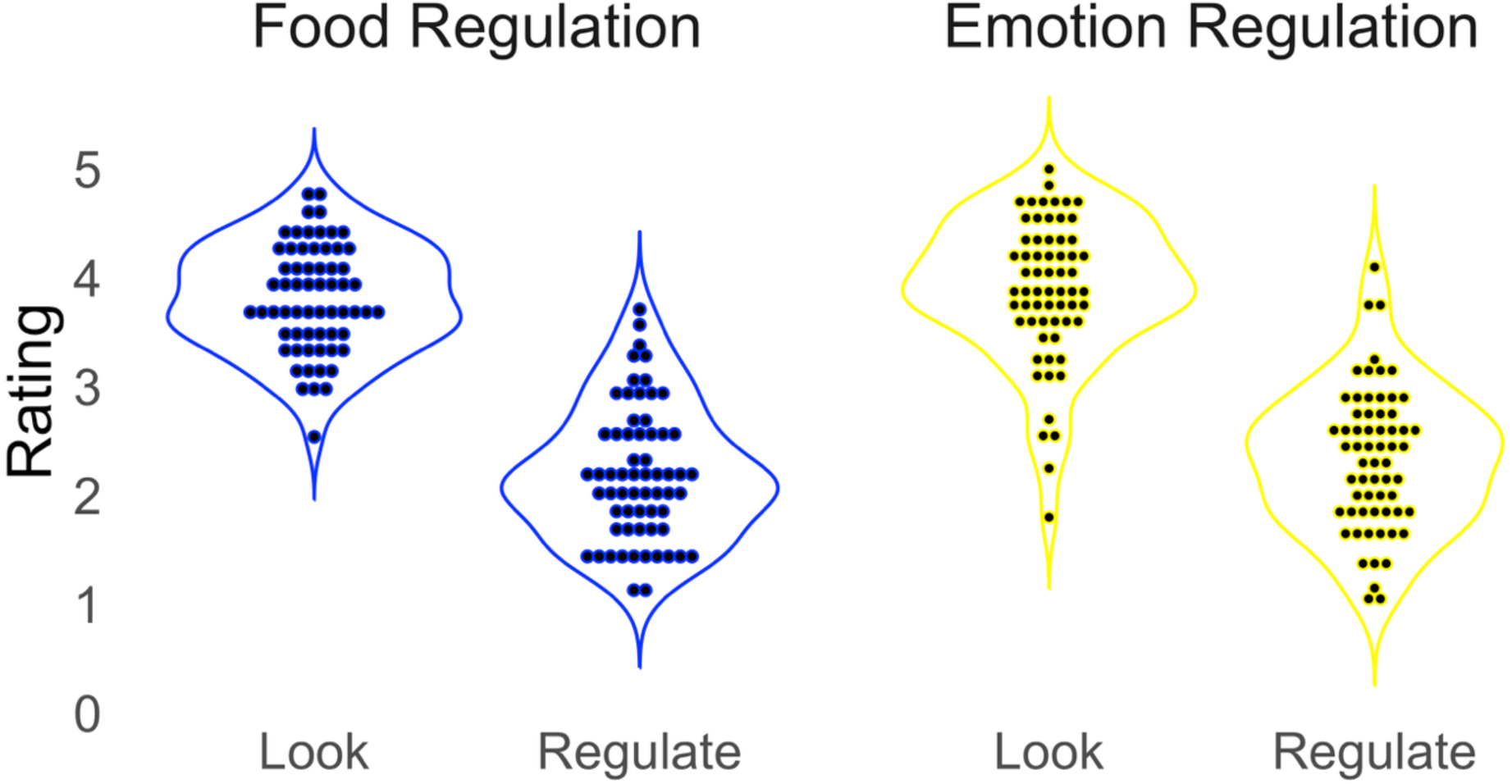
Ratings of food cravings for the FR task and ratings of negative affect for the ER task. Ratings on “LOOK” trials were higher than “DECREASE” (Regulate) trials for both FR (*Mlook* = 3.81, *Mdecrease* = 2.21) and ER (*Mlook* = 3.88, *Mdecrase* = 2.34) tasks, with higher ratings corresponding to higher desire that participants had to eating the food (FR) or feeling more negative while viewing the images (ER). *** = p <.001

We operationalized regulation success in each task as the difference in ratings for look minus regulate trials (FR: *M* =1.59, *SD* = 0.90, *range* = 0.08, −3.3; ER: *M* =1.53, *SD* = 0.85, *range* = 0.5, −3.3). There was a strong positive relationship for regulation success between the FR task and ER task (rho= 0.69, rho^2^ =0.47, p = <0.001, Figure 3).

**Figure 3:**
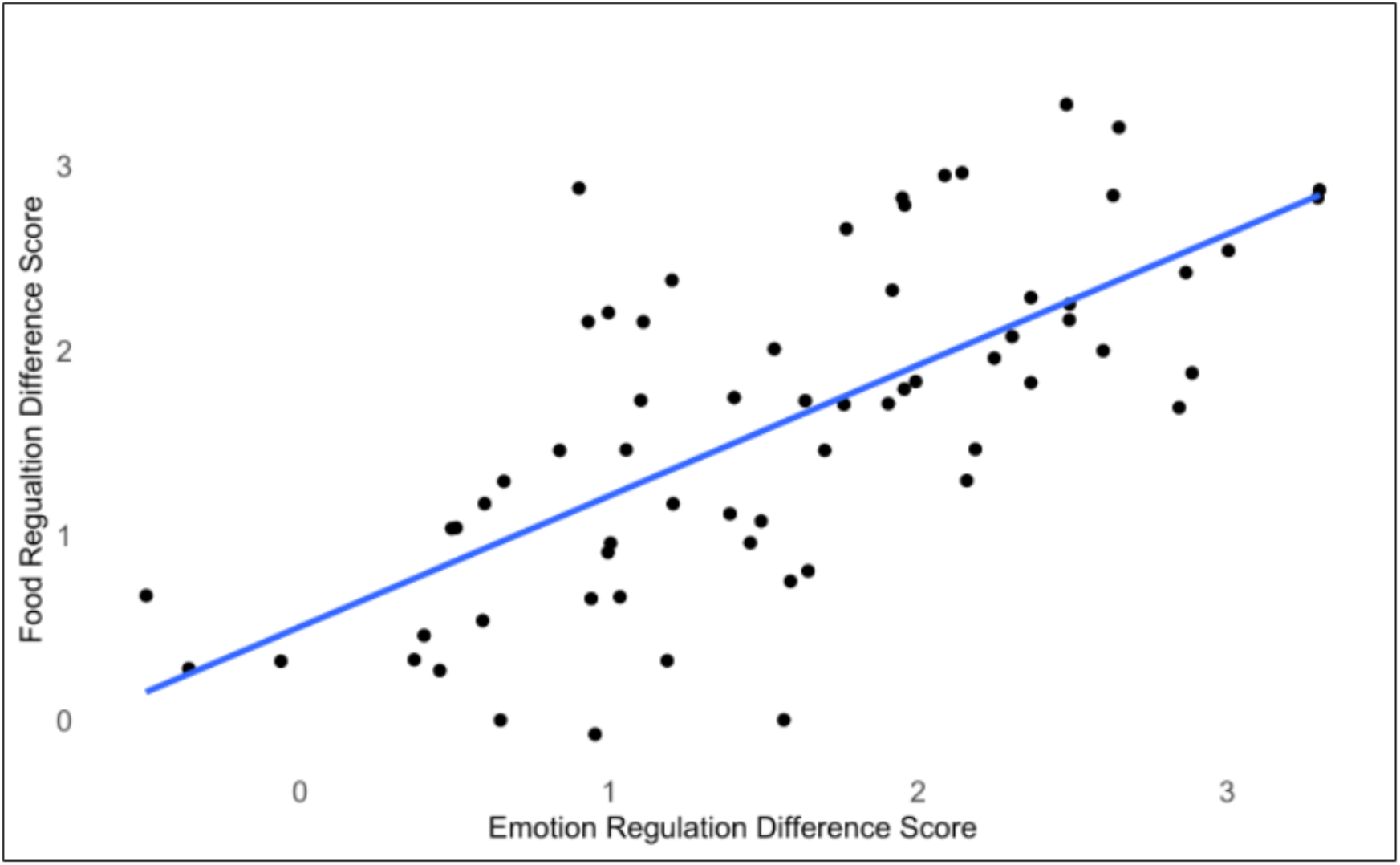
There was a strong positive relationship between performance on emotion regulation and food regulation tasks (rho = 0.69, rho^2^ =0.47, *p* = <0.001). Difference score was calculated by subtracting look - regulate condition.

### fMRI results

#### Domain-general regulation

We conducted a 2 (task: FR, ER) x 2 (condition: regulate, imagine/look) ANOVA on the imaging data. There was a main effect for condition, revealing greater activation for regulate than look trials in portions of the right insula, right inferior and middle frontal gyrus, and bilateral inferior parietal lobule, consistent with previous work (Carnell et al., 2012; Kumar et al., 2016; Li et al., 2021) (Figure 4 and Table 1).

**Figure 4:**
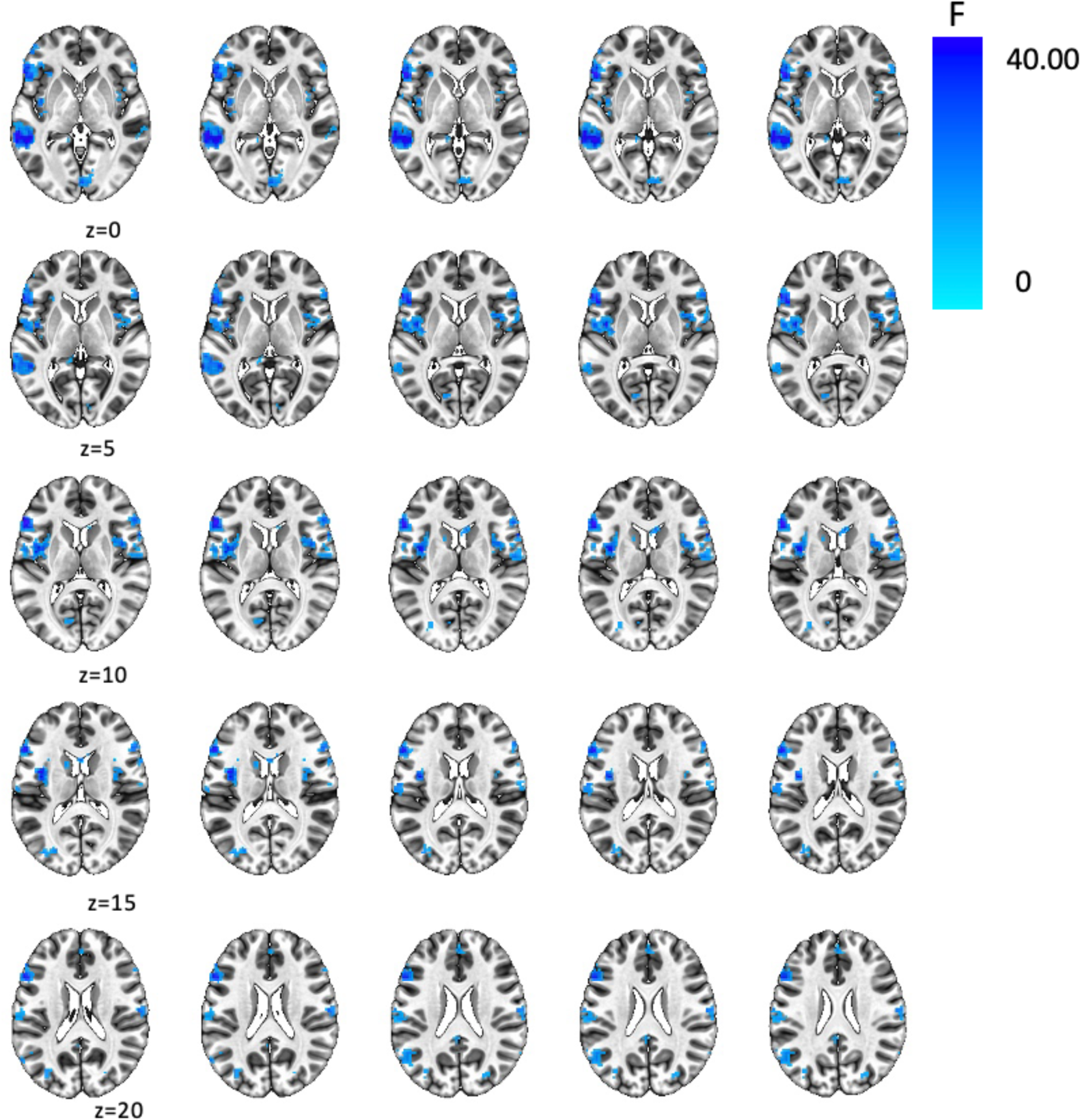
Domain-general (main effect) for food regulation and emotion regulation tasks.

**Table 1:**
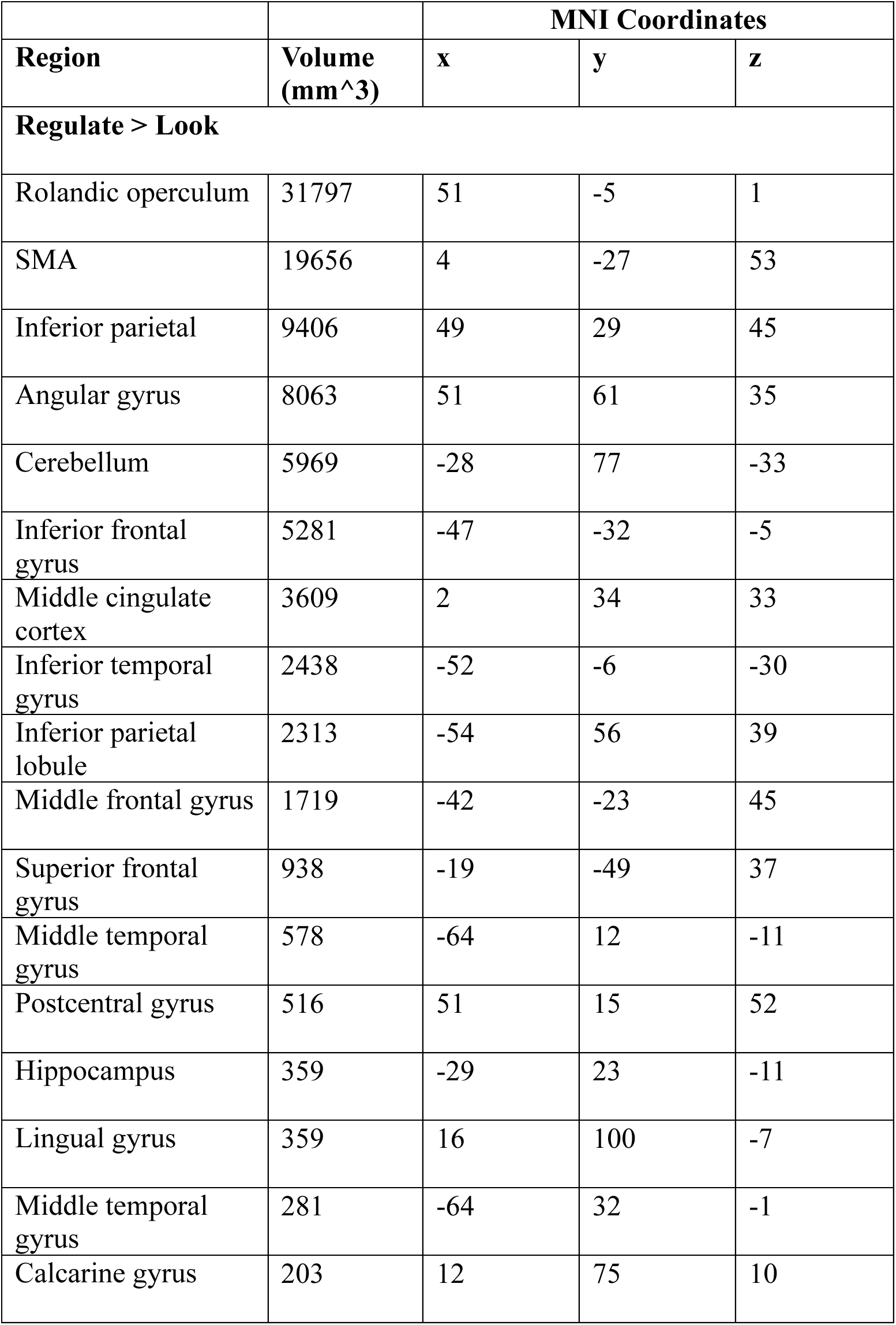
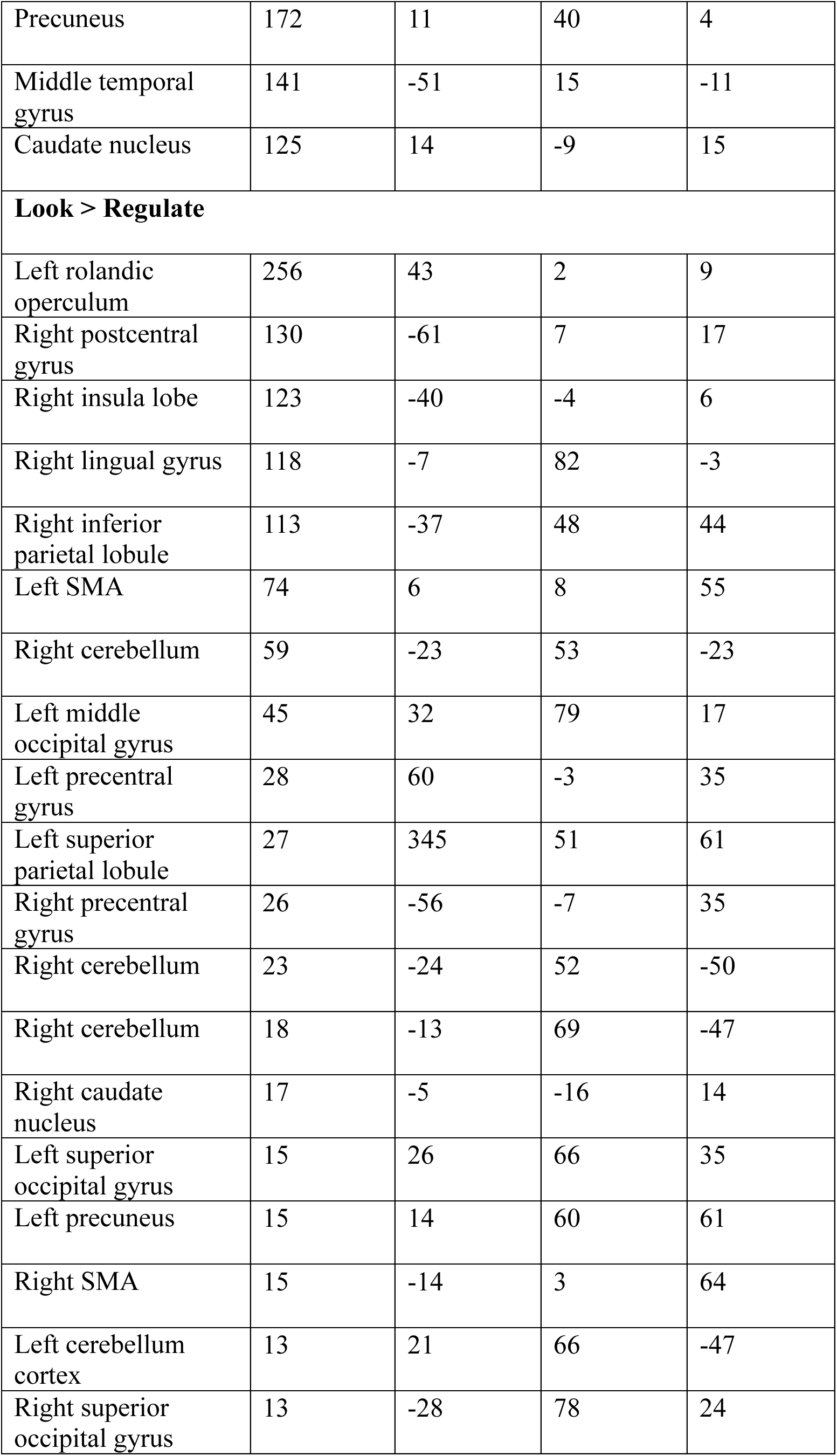
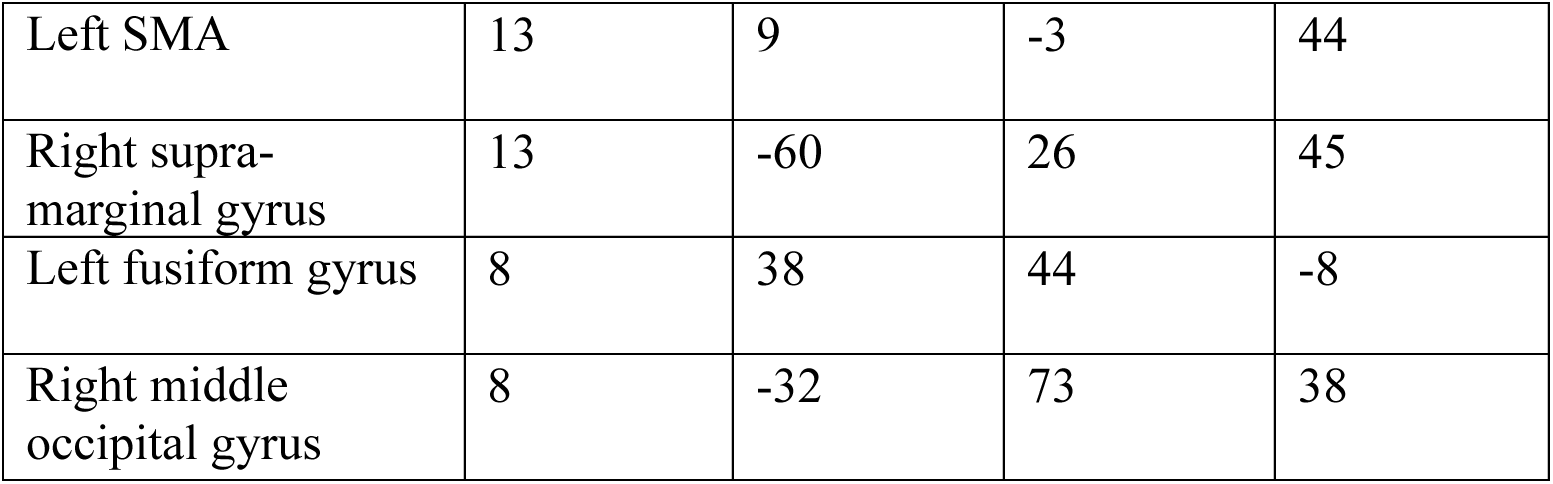
Significant clusters associated with domain-general regulation (main effect) for food regulation and emotion regulation tasks. Note: MNI coordinates are listed as x = RL, y = AP, z = IS (center of mass)

|  |  | MNI Coordinates |  |  |
| --- | --- | --- | --- | --- |
| Region | Volume<br>(mm<sup>3</sup>) | x | y | z |
| **Regulate > Look** |  |  |  |  |
| Rolandic operculum | 31797 | 51 | -5 | 1 |
| SMA | 19656 | 4 | -27 | 53 |
| Inferior parietal | 9406 | 49 | 29 | 45 |
| Angular gyrus | 8063 | 51 | 61 | 35 |
| Cerebellum | 5969 | -28 | 77 | -33 |
| Inferior frontal<br>gyrus | 5281 | -47 | -32 | -5 |
| Middle cingulate<br>cortex | 3609 | 2 | 34 | 33 |
| Inferior temporal<br>gyrus | 2438 | -52 | -6 | -30 |
| Inferior parietal<br>lobule | 2313 | -54 | 56 | 39 |
| Middle frontal gyrus | 1719 | -42 | -23 | 45 |
| Superior frontal<br>gyrus | 938 | -19 | -49 | 37 |
| Middle temporal<br>gyrus | 578 | -64 | 12 | -11 |
| Postcentral gyrus | 516 | 51 | 15 | 52 |
| Hippocampus | 359 | -29 | 23 | -11 |
| Lingual gyrus | 359 | 16 | 100 | -7 |
| Middle temporal<br>gyrus | 281 | -64 | 32 | -1 |
| Calcarine gyrus | 203 | 12 | 75 | 10 |
| Precuneus | 172 | 11 | 40 | 4 |
| Middle temporal gyrus | 141 | -51 | 15 | -11 |
| Caudate nucleus | 125 | 14 | -9 | 15 |
| **Look > Regulate** |  |  |  |  |
| Left rolandic operculum | 256 | 43 | 2 | 9 |
| Right postcentral gyrus | 130 | -61 | 7 | 17 |
| Right insula lobe | 123 | -40 | -4 | 6 |
| Right lingual gyrus | 118 | -7 | 82 | -3 |
| Right inferior parietal lobule | 113 | -37 | 48 | 44 |
| Left SMA | 74 | 6 | 8 | 55 |
| Right cerebellum | 59 | -23 | 53 | -23 |
| Left middle occipital gyrus | 45 | 32 | 79 | 17 |
| Left precentral gyrus | 28 | 60 | -3 | 35 |
| Left superior parietal lobule | 27 | 345 | 51 | 61 |
| Right precentral gyrus | 26 | -56 | -7 | 35 |
| Right cerebellum | 23 | -24 | 52 | -50 |
| Right cerebellum | 18 | -13 | 69 | -47 |
| Right caudate nucleus | 17 | -5 | -16 | 14 |
| Left superior occipital gyrus | 15 | 26 | 66 | 35 |
| Left precuneus | 15 | 14 | 60 | 61 |
| Right SMA | 15 | -14 | 3 | 64 |
| Left cerebellum cortex | 13 | 21 | 66 | -47 |
| Right superior occipital gyrus | 13 | -28 | 78 | 24 |
| Left SMA | 13 | 9 | -3 | 44 |
| Right supra-marginal gyrus | 13 | -60 | 26 | 45 |
| Left fusiform gyrus | 8 | 38 | 44 | -8 |
| Right middle occipital gyrus | 8 | -32 | 73 | 38 |

#### Domain-specific regulation

In addition to the main effect of condition, there was also a significant task x condition interaction. Post-hoc tests of the interaction were conducted by subtracting the imagine/look condition from the regulate condition for each task (Figure 5 and Table 2). Interaction clusters that showed the greatest differential activation for FR > ER included the posterior insula and the right inferior frontal gyrus (Hollmann et al., 2012; Gerosa et al., 2023), whereas the reverse (ER > FR) was evident in the left posterior cingulate cortex and right superior frontal gyrus, consistent with previous research (Goldin et al., 2008; Grecucci et al., 2013) (Figure 4).

**Figure 5:**
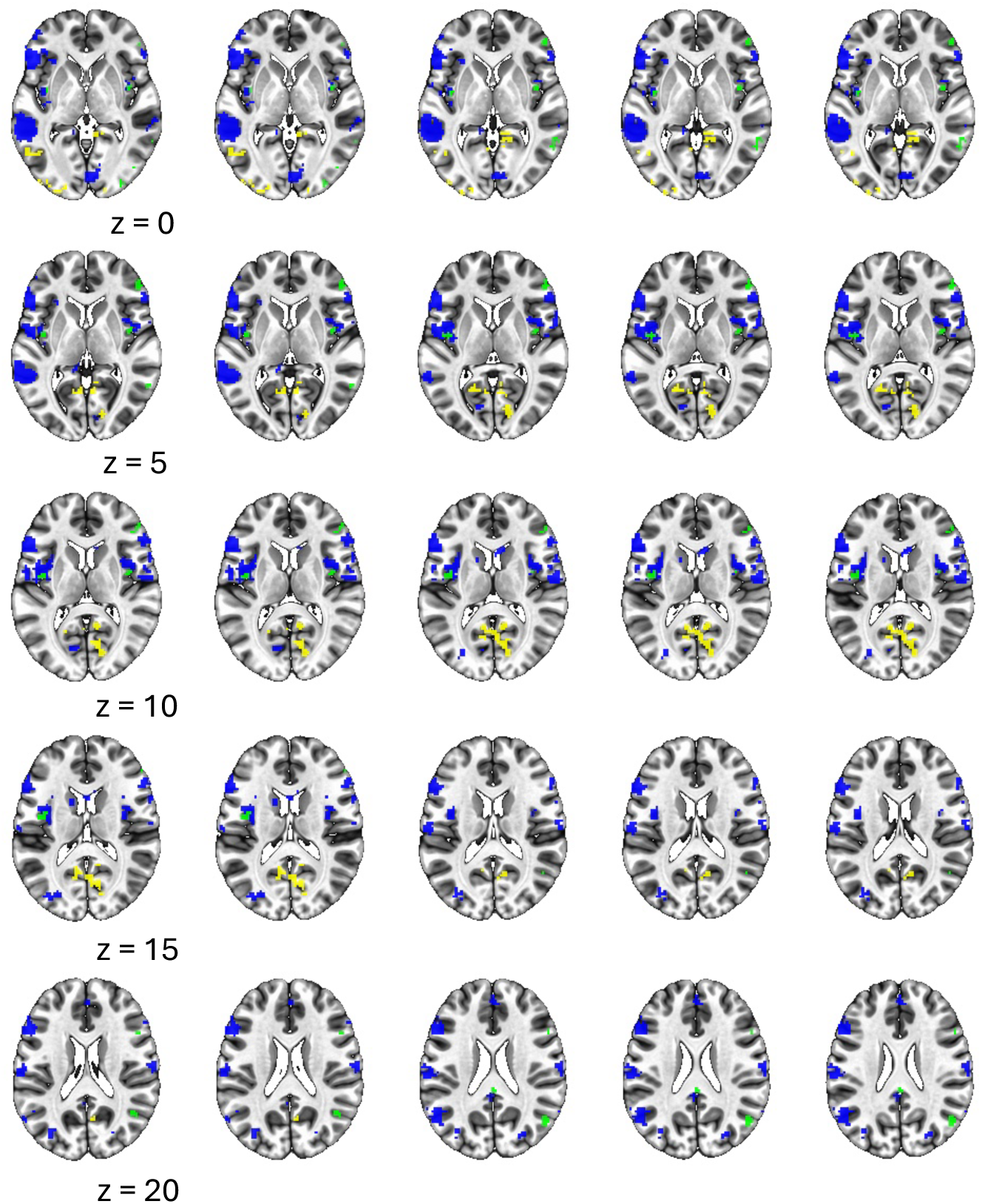
Regions that show greater differential regulation for domain-specific emotion regulation (yellow), domain-specific food regulation (blue), and the domain-general regulation main effect with no interaction (green).

**Table 2:** Domain-specific significant clusters associated with the task x condition interaction. Note: MNI coordinates are listed as x = RL, y = AP, z = IS (center of mass)

|  |  | MNI Coordinates |  |  |
| --- | --- | --- | --- | --- |
| Region | Volume<br>(mm<sup>3</sup>) | x | y | z |
| **Emotion > Food** |  |  |  |  |
| Inferior occipital gyrus | 4578 | 35 | 84 | -6 |
| Calcarine gyrus | 2484 | -7 | 65 | 11 |
| Cerebellum | 1609 | 33 | 50 | -18 |
| Superior frontal gyrus | 406 | -23 | -34 | 47 |
| Calcarine gyrus | 328 | 12 | 59 | 7 |
| Middle orbital gyrus | 250 | -38 | -45 | -10 |
| Superior frontal gyrus | 219 | -25 | -15 | 51 |
| Hippocampus | 203 | 23 | 26 | -7 |
| Superior frontal gyrus | 188 | -21 | -27 | 58 |
| Hippocampus | 156 | -22 | 26 | -6 |
| Lingual gyrus | 156 | -18 | 96 | -2 |
| Cerebellum | 141 | 14 | 45 | -48 |
| **Food > Emotion** |  |  |  |  |
| Middle cingulate cortex | 1641 | 1 | 25 | 30 |
| Cerebellum | 703 | -26 | 40 | -18 |
| Rolandic operculum | 547 | 41 | 5 | 10 |
| Inferior frontal gyrus | 500 | -49 | -41 | 8 |
| Inferior occipital gyrus | 469 | -32 | 92 | -4 |
| Cuneus | 453 | 10 | 74 | 38 |
| Middle temporal gyrus | 406 | -48 | 64 | 23 |
| Insula lobe | 359 | -40 | 1 | 5 |
| Middle temporal gyrus | 203 | -59 | 54 | 3 |
| Postcentral gyrus | 188 | -64 | 13 | 29 |
| Postcentral gyrus | 172 | -53 | 1 | 30 |
| Cuneus | 172 | -13 | 68 | 38 |
| Inferior frontal gyrus | 156 | 45 | -25 | -6 |
| Cerebellum | 141 | -28 | 78 | -36 |
| Cerebellum | 141 | -21 | 87 | -30 |
| Inferior occipital gyrus | 141 | -46 | 77 | -3 |
| Posterior cingulate cortex | 140 | 7 | 43 | 26 |
| Insula lobe | 125 | 41 | -1 | -8 |
| Inferior frontal gyrus | 125 | -52 | -16 | 24 |
| Angular gyrus | 125 | 38 | 68 | 49 |

#### Brain-behavior relationships

We examined the relationship between regulation success and domain-specific brain activation by correlating the difference scores of look > regulate related activity in the interaction clusters with our measure of regulation success (i.e., the difference in ratings between look and regulate for each respective task). There were no statistically significant relationships between regulation success and domain-specific brain activation. We also examined the relationship between regulation success and domain-general brain activation by correlating the difference scores of look > regulate related activity in regions showing a main effect for regulation, excluding voxels showing an interaction (green regions in Figure 5). There were no statistically significant relationships between regulation success and domain-general brain activation.

## Discussion

The present study used a within-subjects design to identify overlapping and distinct neural mechanisms for cognitive reappraisal in the context of FR and ER. There was domain-general activation in the right insula, right inferior and middle frontal gyrus, and bilateral inferior parietal lobule. There was domain-specific activation for the FR task in the posterior insula and for the ER task in the left posterior cingulate cortex and right superior frontal gyrus. Although previous studies have investigated the use of cognitive reappraisal in FR and ER separately and the present results are consistent with these findings, the present study makes novel contributions by identifying overlapping and unique regulatory mechanisms across these tasks using a within-subjects design. This research provides important insight into domain-general and domain-specific effects of regulation that could theoretically inform interventions for emotion and food dysregulation.

In both the FR and ER tasks, participants had decreased ratings in the “regulate” condition relative to the “look” condition, demonstrating successful ability to use the cognitive reappraisal strategy. Additionally, there was a strong relationship between regulation success on both tasks such that participants who were able to effectively regulate emotion cues were also able to effectively regulate food cues. However, there was no significant relationship between regulation success and brain activation (i.e., activation did not vary as a function of how successfully participants performed the respective tasks). Due to task performance on one task explaining about 47% of the variance in performance on the second task, it’s important to consider both domain-general and domain-specific cognitive control since these results indicate that, while there may be some overlap between regulatory cognitive processes (i.e., domain-general), there are also differences (i.e., domain-specific). This set of findings provides further evidence that a core set of brain regions are associated with cognitive control across domains, and there are also distinct brain regions involved in regulation within specific task domains.

Our results are consistent with previous between-subjects studies of cognitive reappraisal that show overlapping neural activation for FR and ER. The findings that the right inferior and middle frontal gyrus and the bilateral inferior parietal lobe are involved in domain-general regulation are consistent with previous findings that brain networks involving the PFC (e.g., fronto-parietal and cingulo-opercular networks) are critical for regulatory behavior such as inhibition and attentional control (Chambers et al., 2009; Friedman & Robbins, 2021; Menon & Esposito, 2021; Zanto & Gazzaley, 2014), and that the parietal cortex is involved in initiating regulatory behavior (Chen et al., 2022). Activation in control brain networks appear to be critical in regulatory processes, regardless of whether participants are regulating responses to food or negative emotions. This finding suggests that there is an overlapping mechanism supporting regulation that is required across various domains or instances of reappraisal.

The evidence for an underlying regulation mechanism has potentially important implications for intervention. Indeed, cognitive control is critical to being able to regulate one’s food cravings and negative emotions. This study highlights similarities in these two regulatory processes suggesting that one intervention could potentially target and improve both food- and emotion-related outcomes. For example, current research has shown that 20-30 minutes of moderate-intensity aerobic exercise, which increases functional connectivity among control-related brain regions (e.g., left dorsolateral PFC, posterior cingulate cortex, and the right inferior parietal cortex), may improve cognitive control across the lifespan (Themanson, Pontifex & Hillman, 2008, Hillman et al., 2009, Weng et al., 2017). In the present study, the dorsolateral PFC and posterior cingulate cortex showed domain-specific activation for FR and ER, respectively. Given the role of cognitive control in regulatory behavior, and the overlap of activation found in studies of aerobic exercise and of FR and ER, aerobic exercise is one area of potential future research for intervention to improve domain-general regulation.

Above and beyond the overlapping neural mechanism, differences were also observed between these two tasks, suggesting some aspects of regulation are domain-specific. Consistent with our findings, previous neuroimaging studies have described activation in brain regions associated with reward processing and the sensory perception of food-regulated stimuli, such as the insula, during the cognitive reappraisal of food (Frank et al., 2013; Kumar et al., 2016). We also observed greater differential activation in the left posterior cingulate cortex for emotion regulation, consistent with literature showing the involvement of the posterior cingulate cortex to the processing of emotions (Rolls, 2019), autobiographical memory (Auger & Maguire, 2013; Leech & Sharp, 2014), altering responses based on task demands (Pearson et al., 2011), and self-regulatory behavior (Kirlic et al., 2022).

Some limitations of the study should be noted. First, average participant BMI fell within the healthy weight range, which makes it unclear if there would be unique regulatory behavior responses (particularly for FR) for participants that fell in the underweight, overweight, or obese BMI categorizations. Second, although participants were instructed to either passively look at the images or use a cognitive reappraisal technique while viewing images in both tasks, the instructions for the FR task specifically instructed participants to reappraise by focusing on the health consequences of eating the desirable food, whereas the ER task instructions were more open-ended about how to apply reappraisal. The greater specificity in the FR instructions may have influenced the differences in brain mechanisms between the two tasks. A future study could either narrow the ER instructions to a single approach or broaden the FR task instructions to cognitively reappraise the food stimuli to make it less appetizing (rather than relating it to health consequences of eating the food). Third, although we did not find a statistically significant relationship between regulatory success and brain activation, the literature suggests that these behaviors are related to success during the ER task. One possible explanation for the lack of replicating the behavior-brain relationship is due to our limited sample size. Lastly, participants were undergraduate students enrolled in a psychology course, so it is unclear to what extent these findings would generalize to other samples.

The current study builds on the existing literature by identifying overlapping brain regions that are associated with cognitive reappraisal during FR and ER tasks, and brain regions that are differentially associated with these two regulatory processes. The existing literature has previously examined these regulatory processes separately, and the current study results expand on these findings in a within-subjects context, paving the way for future research identifying and testing interventions for related mental and physical health outcomes. Future research should determine the extent to which these domain-general and domain-specific mechanisms are also evident in clinically relevant populations (e.g., individuals suffering from food and/or emotion dysregulation, such as those who are overweight or obese, or those presenting with mental health problems).

## Author CRediT Statement

J.L., C.S., M.N., T.N., and D.S. contributed to the conceptualization of the project; T.N. acquired funding and resources for the project, and oversaw the investigation process and data collection; J.L. and C.T. contributed to the data curation; J.L. performed the formal analysis, visualization of data, and took the lead in writing the manuscript; J.L., T.N., and D.S. contributed to the methodology; D.S. provided supervision to all aspects of manuscript preparation (i.e., data analysis interpretation and revisions for original draft prior to sending to other co-authors); M.N. and D.S. provided software (i.e., E-Prime tasks) for the project; C.S., M.N., C.T., T.N., and D.S. reviewed and edited the manuscript.

## Data Availability Statement

The data for this project cannot be shared publicly due to participants not consenting to having their data shared at the time of study participation. Please contact the corresponding author, Douglas Schultz, with additional questions on data availability.

## Acknowledgements

This research was supported by the National Institute of General Medical Sciences (U54GM115458) and the National Institute of Diabetes and Digestive and Kidney Diseases (R01DK125651, F31DK122636) of the National Institutes of Health (NIH). The content is solely the responsibility of the authors and does not necessarily represent the official views of the NIH.

